# A roadmap for a quantitative ecosystem-based environmental impact assessment

**DOI:** 10.1101/080242

**Authors:** J. Coston-Guarini, J-M Guarini, J. Edmunds, Shawn Hinz, Jeff Wilson, L. Chauvaud

**Affiliations:** The Entangled Bank Laboratory, Banyuls-sur-Mer, 66650 France; Ecole Doctorale des Sciences de la Mer, UBO, CNRS, UMR 6539-LEMAR IUEM Rue Dumont d’Urville Plouzané, 29280 France; Laboratoire International Associé ‘BeBEST’, UBO, Rue Dumont d’Urville Plouzané, 29280 France; LimOce Environmental Consulting, Ltd. 47 Hamilton Sq. Birkenhead, CH41 5AR UK; Gravity Environmental Consulting, Fall City, WA, 98024 USA; CNRS, UMR 6539-LEMAR IUEM Rue Dumont d’Urville Plouzané, 29280 France

**Keywords:** Environmental Impact Assessment, ecosystem, drivers of change, modelling, socio-ecological system

## Abstract

A new roadmap for quantitative methodologies of Environmental Impact Assessment (EIA) is proposed, using an ecosystem-based approach. EIA recommendations are currently based on case-by-case rankings, distant from statistical methodologies, and based on ecological ideas that lack proof of generality or predictive capacities. These qualitative approaches ignore process dynamics, scales of variations and interdependencies and are unable to address societal demands to link socio-economic and ecological processes (*e.g.* population dynamics). We propose to re-focus EIA around the systemic formulation of interactions between organisms (organized in populations and communities) and their environments but inserted within a strict statistical framework. A systemic formulation allows scenarios to be built that simulate impacts on chosen receptors. To illustrate the approach, we design a minimum ecosystem model that demonstrates non-trivial effects and complex responses to environmental changes. We suggest further that an Ecosystem-Based EIA - in which the socio-economic system is an evolving driver of the ecological one - is more promising than a socio-economic-ecological system where all variables are treated as equal. This refocuses the debate on cause-and-effect, processes, identification of essential portable variables, and a potential for quantitative comparisons between projects, which is important in cumulative effects determinations.

## INTRODUCTION

When the USGS hydrologist and geomorphologist Luna Leopold (1915-2006) and his two co-authors published a system for environmental assessment in 1971 (Leopold *et al.*, 1971), they could not have foreseen that 50 years later, their report would be at the origin of a global industry (Morgan, 2012; Pope *et al.*, 2013). Leopold *et al.* produced their brief document at the request of the US Department of the Interior after the National Environmental Policy Act (NEPA) created a legal obligation for federally funded projects to assess impact. In the year following the passage into law, the scientific community was quick to point out the absence of any accepted protocol for either the content of the document or its evaluation (see characterisation in Gillette, 1971). In response, Leopold *et al.* describes a preliminary approach, with a simple decision-tree like diagram (Figure 1A) relying on structured information tables. These tables of variables and qualities, or ‘interaction matrices’, are intended to enforce production of uniform, comparable descriptions, while requiring only a minimum of technical knowledge from the user.

**Figure 1.**
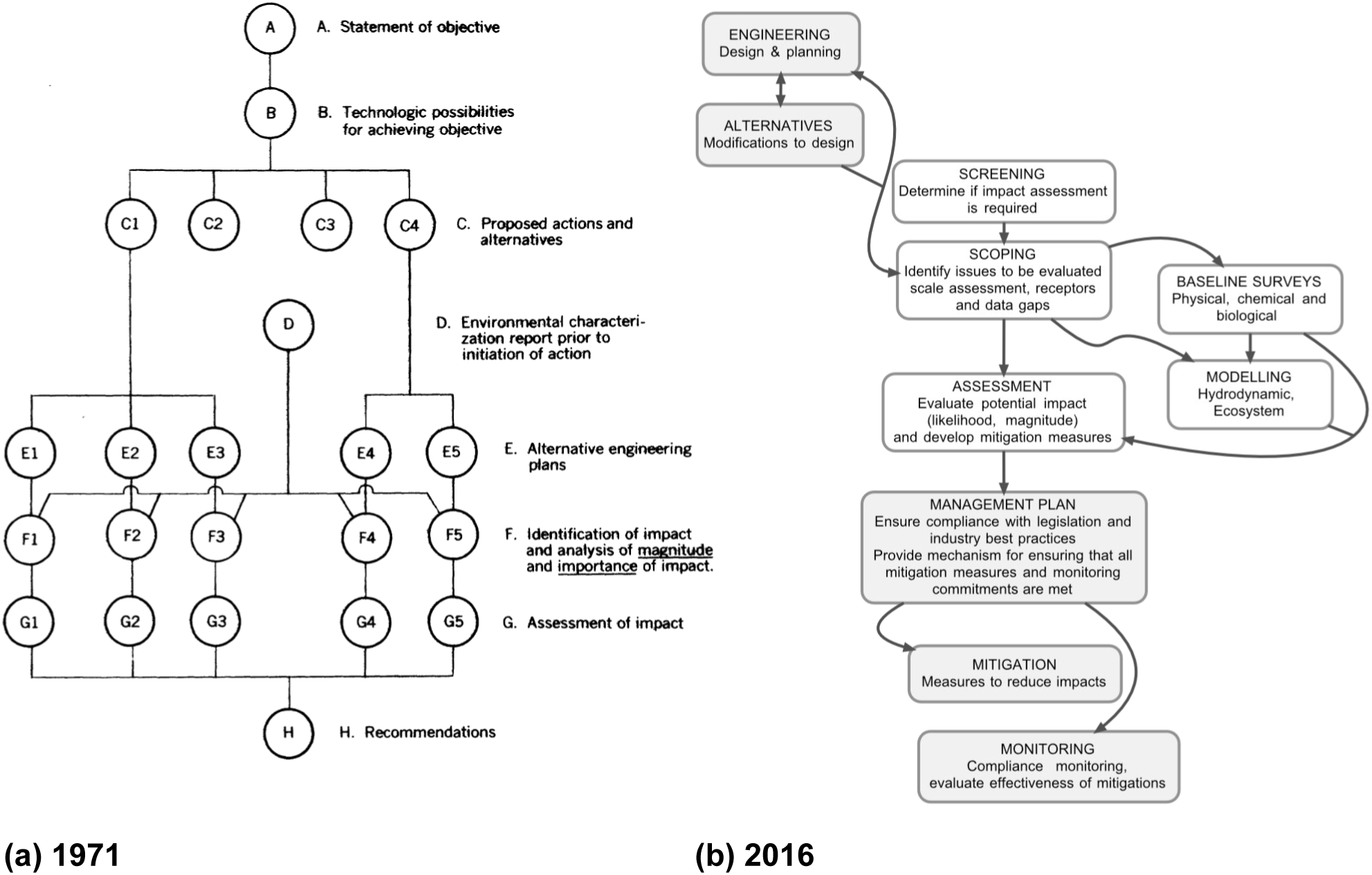
Environmental impact assessment, then and now. (a) The original flow chart as it appeared in Leopold *et al.* 1971. This chart responds to a specific request by the US Department of the Interior to propose a system that would structure information in EI documents. The original figure is captioned: “Evaluating the environmental impact of an action program or proposal is a late step in a series of events which can be outlined in the following manner.” (b) Example of a flow chart used by consultants today in offshore projects. Important changes include: the addition monitoring and the possibility of using modeling. Steps external to the core EIA steps are in grey. Redrawn after Edmunds *et al.* 2016.

Impact inference rests on a statistical comparison of variables between impacted and non-impacted sites, but assessing an impact is understood to include value-based judgements about quality and importance (Leopold *et al.*, 1971) linked with attitudes held about the environment (Buttel and Flinn, 1976; Lawrence, 1997; Toro *et al.*, 2013). These judgements, often made *a priori* (Toro *et al.*, 2013), can conflict with the necessity to reach a legal standard of proof (Goodstein, 2011) when projects are contested. EIAs therefore embody a compromise between technical descriptions of the expected magnitude of an impact on a receptor and managerial recommendations about how to avoid that receptors exceed acceptable values, or mitigate, identified impacts (Lawrence, 1997; Cashmore *et al.*, 2010; Barker and Jones, 2013). By 1971, under pressure to move development projects forward (Gillette, 1971), the EIA process became institutionalised as a qualitative exercise focussed on collecting documentation about a project site supported by individuals’ professional expertise, without requiring quantitative evaluations to back up statements (Lawrence, 1997; Cashmore *et al.*, 2010; Morgan, 2012; Toro *et al.*, 2013). Hence EIAs today still strongly resemble the preliminary instructions given by Leopold *et al.* (Figure 1B). Consequently, review articles, such as that of Barker and Jones (2013) on offshore EIAs in the UK, often report strong criticisms of the quality of environmental impact documents as being “driven by compliance rather than best practice”.

Over the past decade, technologically sophisticated monitoring tools and baseline surveys have been integrated (*e.g.* Figure 1B, “Modelling”; Payraudeau and van der Werf, 2005; Nouri *et al.*, 2009) on a discretionary basis because they contribute to risk management of sensitive receptors as well as to new dynamic features like the “Life Cycle Assessment” of a project (Židonienė and Kruopienė, 2015). These changes suggest that EIA is poised to incorporate quantitative frameworks.

Inspired by the application of ecosystem-based management frameworks in fisheries (Smith *et al.*, 2007; Jacobsen *et al.*, 2016), and by the generalisation of modelling and statistical tools in ecological and environmental sciences, we describe in this article how the objective of a quantitative, ecosystem-based EIA could be achieved. We first examine briefly the awareness of impact and analytical approaches that exist to quantify this within ecological sciences. We then propose a quantitative reference framework linking statistical impact assessment to ecosystem functioning and discuss how the modelling approach may be used to provide reasonable predictions of different categories of impact. Finally, we explore how our ecological system will behave when socio-economic “drivers of change” (UNEP, 2005) are implemented. By imposing socio-economic factors as drivers (instead of as variables of a large integrated system), we show that different types of consequences can occur, which are not represented by classical feedbacks. For example, this permits the life cycle of the project to be described as a driver of the dynamic of the impacted system, or the explicit implementation of cumulative effects scenarios.

### Awareness of environmental impact in the past

There is a long written record of the awareness that human activities affect the environment. Texts of 19^th^ century naturalists commonly contain remarks about the disappearance of animals and plants attributed to human activities; some are quite detailed, like George P. Marsh’s quasi-catalogue of the ways “physical geography” (natural environments) has been altered by development (Marsh, 1865). Most are ancillary comments to make rhetorical points, rather than scientific observations, like this quote from the marine zoologist Henri de Lacaze-Duthiers (1821-1901) (de Lacaze-Duthiers, 1881: 576-577):

*“Ainsi, lorsque sera crée la nouvelle darse, qui n'a d'autre but que d'augmenter le mouvement du port, que deviendront les localités tranquilles où la faune était si riche ? Resteront-elles les mêmes ? l’eau ne se renouvelant pas, n'aura-t-elle pas le triste sort de celle des ports de Marseille, si le commerce et les arrivages prennent de grandes proportions ?*

*“Le mouvement du port augmente tous les jours. Les constructions des darses projetées ne modifieront-elles pas les conditions favorables actuelles ? On doit se demander encore si l'eau conservera son admirable pureté quand le nombre des bâtiments aura augmenté dans les proportions considérables que tout fait prévoir.*

*“Port-Vendres ne peut évidemment que se modifier profondément dans l’avenir, et cela tout à l’avantage du commerce, c’est-à-dire au détriment de la pureté, de la tranquillité de l’eau et du développement des animaux.*

*A Banyuls, il n’y a aucune crainte à avoir de ce côté.”*

When he wrote this, Lacaze-Duthiers had been lobbying for more than a decade for the creation of a network of marine stations in France. His text justifies why he chose a village without a port, instead of one with a thriving port. His reasoning is that economic development causes increases in buildings, docks, boat traffic, that damages the “tranquillity”, “water purity”, and the “favourable conditions for development of fauna”. While he acknowledges this is a gain for local commercial interests, it is also at the expense of faunal richness, and he predicts this will lead to the “sad situation of the port of Marseille”. Lacaze-Duthiers feels this degradation should be a legal issue or a civil responsibility (as “*au détriment de*” indicates a legal context). The attitude and awareness of Lacaze-Duthiers are symptomatic of ambiguities about the environment (Nature) and the place of humans in it, that are also at the core of EIA (Cashmore, 2004; Wood, 2008; Morgan, 2012; Toro *et al.*, 2012). These political conflicts between a desire to preserve the natural world and its own functioning, and the desire to use, exploit, order and control parts of it are the main issues of impact assessment (Cashmore *et al.*, 2010).

### Path to reconciliation

What changed in the latter half of the 20^th^ century is that managers, regulators and stakeholders need to document and quantify impacts as well as their associated costs. However, important, historical contingencies complicated the development of quantitative tools for environmental impact. Ecosystem science, which pre-dates EIA by several decades, describes ecosystem functioning in terms of energy and mass flows (*e.g.* Odum, 1957) and the distribution of species is understood with respect to how well the ‘conditions of existence’ of a population are met and maintained (*e.g.* Gause, 1934; Ryabov and Blasius, 2011; Adler *et al.*, 2013). These approaches use paradigms from biology, physics and chemistry to describe functions and quantify fluxes. Consequently, ecosystem science was not concerned with characterising environmental quality, but determining when conditions of existence were met within dynamic, interacting systems. By the 1970s when EIA practice emerged, ecological research was busy with adaptation and community succession (Odum, 1969; McIntosh, 1985), while the concepts of environmental quality and impact were being defined under a “political imperative, not a scientific background” (Cashmore, 2004: 404) using static components like receptors and indices.

Today, several very different, co-existing strategies exist with regards to environmental management and conservation: ecosystem functioning (*e.g.* Moreno-Mateos *et al.*, 2012; Peterson *et al.*, 2009), ecosystem services and markets analysis (*e.g.* Beaumont *et al.*, 2008; Gómez-Baggethun *et al.*, 2010), and environmental impact. In this context, knitting together sociological and ecological frameworks has emerged as a very active area of interdisciplinary research (Binder *et al.*, 2013). An important theme has been to re-conceptualise environmental dynamics from an anthropogenic perspective to counter a perception that human activities have been excluded from ecological studies (Berkes and Folke, 1998; Tzanopoulos *et al.*, 2013). While this is clearly an unfair characterization (the classic introductory American text on ecology is entitled “Ecology: The link between the natural the social sciences”; Odum, 1975), we do recognize that, historically, ecological sciences have often ignored human behaviours and attitudes in ecosystem studies, despite numerous appeals (Odum, 1977; McIntosh, 1985; Berkes and Folke, 1998). Inspired by the criticisms of Lawrence (1997) about EIA and the challenge of working between both sociological and ecological systems (Rissman and Gillon, 2016), we propose a quantitative basis for systems-based impact assessment. Our goal is to renew the understanding of impact in terms of the interactions and functions attributable to ecosystem processes, integrating the full dynamics of physical and biological processes, while allowing for effective evaluation of socio-economic dynamic alternatives within the modelling framework.

## METHODOLOGY

### Receptors

Assuming that the screening process has already demonstrated the requirement to perform an EIA for a given project, scoping identifies the receptors and the spatio-temporal scale of the study. Receptors are represented by variables being impacted by the project implementation. Receptors are determined by the experts in charge of the EIA. Their qualifications as receptors imply that they will be impacted and this cannot be questionable. In other words, what is called "testing" impact is statistically limited to a process of deciding if the observation data corresponding to samples of the receptor variables permits an impact to be detected. In no case should the selection of a receptor be made with the objective to decide *if there is* an impact or not. By definition, receptors are selected because they are sensitive to the impact. However, all declared receptor variables also represent objects of ecology and can be inserted into an ecosystem framework. These two points will now be reviewed in more detail, establishing an explicit link between them.

### Statistical rationale for impact assessment detection

Impact assessment relies on statistical comparisons of receptor variables in impacted and non-impacted situations. Assuming that the expertise determined the nature of the impact (*i.e.* decreasing or increasing the variable), the impact assessment consists of testing if the absolute difference, Δ, between the non-impacted (µ_0_) and the impacted variable means (µ_I_) is greater than zero (H_1_: Δ = |µ_0_-µ_I_| > 0). Classical testing procedure leads not to accepting H_1_, but to rejecting H_0_ (H_0_: Δ = |µ_0_-µ_I_|= 0). However, the power of the test increases when Δ increases, which means that the more the variable is sensitive, the greater the impact has a chance to be detected.

Ideally, as EIAs start before the project implementation, samples of receptor variables are collected before and, then after the project. We focused on this case even if sampling may also be carried out concurrently for comparing non-impacted and impacted zones. For a receptor variable x, considering two samples of sizes n_0_ (before implementation) and n_I_ (after implementation), the empirical averages are 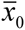 and 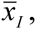 respectively, and their standard deviations are s_0_ and s_I._ The statistics of the test is then:

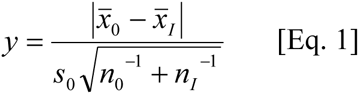

emphasizing the importance of the sample (before implementation), which is used to estimate the ‘baseline’. The dispersion around the average s_0_ has a crucial role in the calculation of y (y decreases when s_0_ increases). Besides the size n_0_ will be fixed when the project is implemented (*i.e.* it is impossible to come back to the non-impacted situation when the project is implemented), while n_I_ can be determined and even modified *a posteriori*.

Under H_1_ (hence when H_0_ is rejected), y is normally distributed, y ~ N(Δ,1), and then it can be centred using:

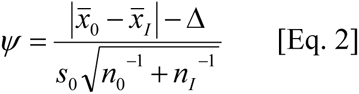

This allows us to state that Ψ follows a Student law. Therefore the test leads to rejection of H_0_ if Ψ is greater than a threshold t_*υ*,*α*_, where *υ* is the number of degrees of freedom (*υ* = n_0_-1) and *α*, the type 1 error (rejecting H_0_ when H_0_ is true), is *α* = proba{Ψ> t_*υ*,*α*_ | Δ=0}). The type 2 error (failing to reject H_0_ when H_0_ is false) is then *β* = proba{Ψ> t_*υ*,*α*_ | Δ>0} and the power of the text is *π*=1-*β*.

As Ψ follows a Student law:

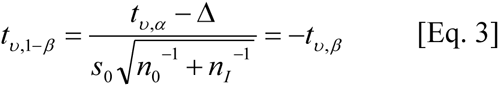

Considering that the baseline is estimated by a sampling performed before implementation, with n_0_ becoming a fixed parameter, the question of detecting significantly the impact then consists of determining two unknown variables Δ and n_I_ by solving two functions:

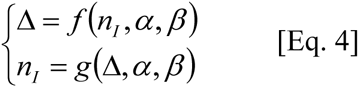

By introducing δ=Δ/µ_0_, the variation Δ relative to the baseline, and 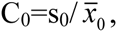, the variation coefficient of the baseline sample, the system to solve is then:

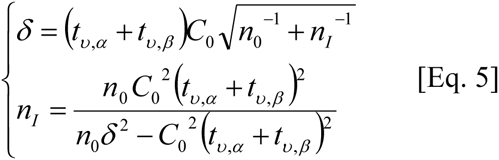

At this point in our development, we can make several remarks about how EIA practices shape the calculation of the impact:

1. The change relative to the baseline (*δ*) is positive if 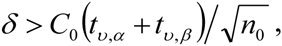 and hence 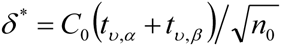 is the detection limit of the receptor variable which can be calculated *a priori* (before impact). δ^*^ is the smallest absolute relative difference that can be characterized, and it depends only on s_0_ and n_0_ and the choice of Type 1 and 2 errors. Therefore, the quality of the expertise, which determines the receptors and the baseline, is a fundamental component of impact assessment.
2. The parametric framework has many constraints (*i.e.* homogeneity and stability of the variance, stability of the baseline …), which have to be ensured, but is very useful for establishing a link with modelling. In particular, µ_0_ and µ_I_, hence *D* and *d*, are descriptors of the states of the impacted ecosystem which can be simulated by calculation from a deterministic model.
3. *A fortiori*, the change relative to the baseline, δ, which depends on the nature of the impact and the temporal scale of the observations, can be determined *a priori* (or plausibly predicted) by the deterministic model. However it implies assuming that the variations which create the dispersion around the trend of the variable are white noises, e_t_ (defined by {E(e_t_) = 0, E(e_t_²) = s_0_, E(e_ti_,e_tj_) = 0}. In this case, the design of the ecosystem becomes particularly important, not only for diagnosing the amplitude of the impact, but also the exact condition of the survey (*i.e.* calculation of n_I_).

### Building an ecosystem model with receptors

Our means to reconcile impact assessment with the theory of ecology is to replace the notion of receptors into a dynamic ecological model (Figure 2A). Receptors are placed in a network of interactions which represent an ecosystem. The “ecosystem” is a system in which the living components will find all conditions for their co-existence in the biotope (abiotic components and interactions that living organisms develop between themselves and with their environment). This classical definition (Tansley, 1935) encounters problems when translated into systemic frameworks. In particular, if the notion of co-existence is often linked to stable equilibrium, there is not one single definition of the notion of stability (Justus, 2008) and the precise nature of the complexity-stability relationship in ecosystems remains unsettled (Jacquet *et al.*, 2016).

**Figure 2.**
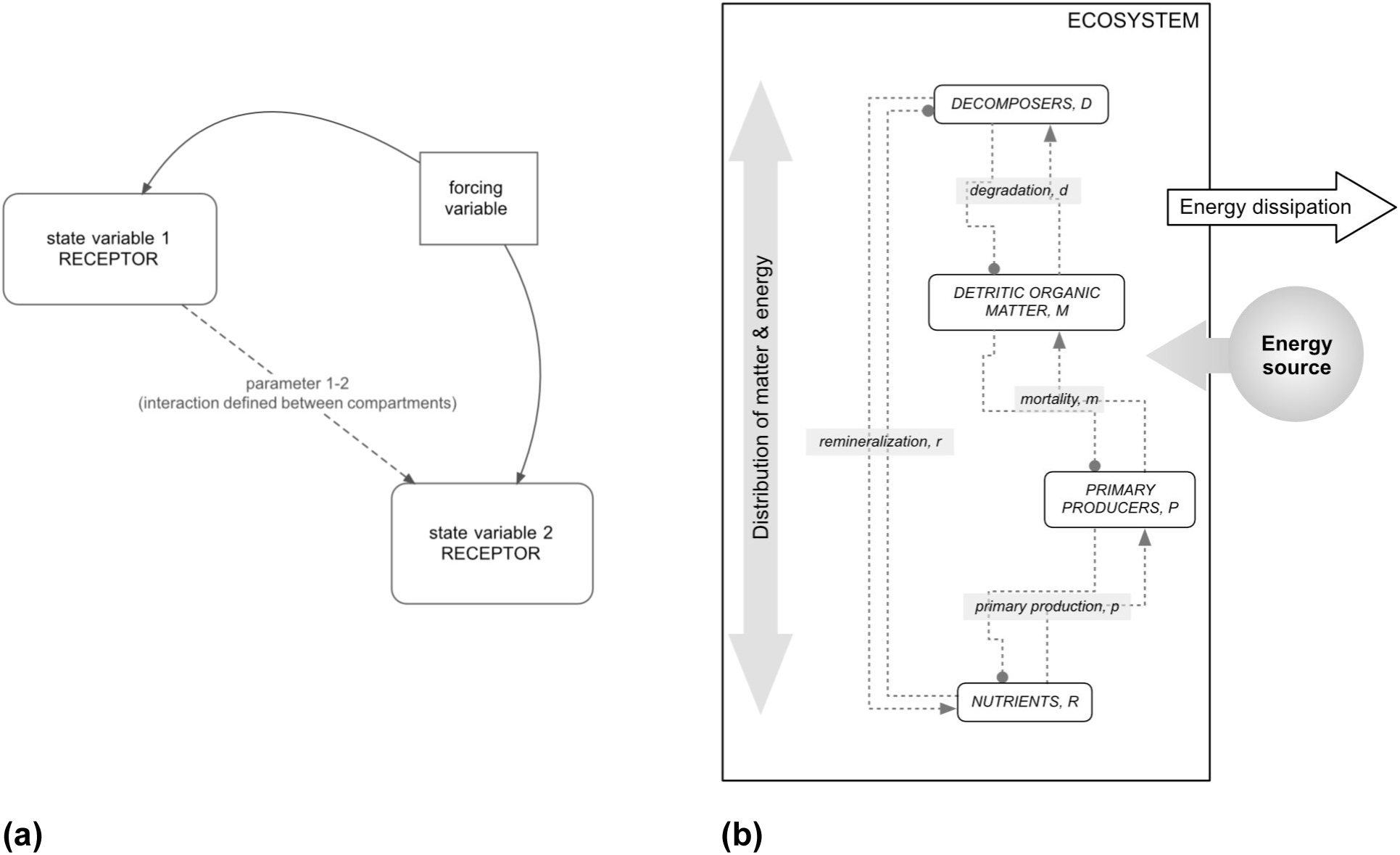
The minimum ecosystem model. (a) The simplest representation of a model in ecology requires two state variables at least one parameter and a ‘forcing’ variable to describe the external forcing by dynamic environmental conditions such as light, temperature, tides. State variables (compartments) are written as a function of the parameters, forcing variables, or other state variables, for a given time interval. Because these vary dynamically they are written as differential equations. Forcing variables are fixed externally, and are not affected by the model calculation of the interaction represented between the two state variables (dashed line). (b) The minimum ecosystem model used in this article is closed in matter but not energy, the energy source is unlimited (forcing variable) and the environment is well-mixed. Feedback interactions between the receptors (state variables) are shown using Levin’s notation, where positive feedback is indicated by arrows and the negative feedback direction is shown by filled circles. Parameter values may be taken from the literature, experiments or field observations.

Even with these caveats, the formulation is useful to explore a system-based EIA. First, stable equilibrium, for a given time scale (from the scale of the project implementation to the of the project life cycle scale) ensures that the baseline would not be subject to drift. Thus, variations will be due to the impact of the project and not by other sources. Secondly, spatial boundaries have to be determined such that the ecosystem has its own dynamics, even if it exchanges matter and energy with other systems. The stable equilibrium is then conditioned by the ecosystem states and not by external forcing factors. This last criterion ensures that the impact can be observable, and not masked by external conditions to the project. At the same time, boundaries are defined by the actual system under investigation and not by the presumed extended area influenced by the project.

For sake of simplicity, we proposed to consider a minimum ecosystem model (Figure 2B). A minimum ecosystem has to ensure the co-existence of two populations: one population accomplishes primary production from inorganic nutrients, and a second degrades detrital matter generated by the first population to recycle nutrients. Hence, there must be four different compartments (pool of nutrients (R), population of primary producers (P), population of decomposers (D) and a pool of detrital organic matter (M)), plus the corresponding four processes linking them, namely, primary production, mortality of primary producers, degradation of detrital organic matter, and remineralization (Figure 2B). Remineralization is linked to the negative regulation of the population of decomposers. Our ecosystem is considered as contained within a well-defined geographic zone (*e.g.* it has a fixed volume), receiving and dissipating energy, but not exchanging matter with the ‘exterior’. The energy source is considered unlimited and not limiting for any of the four biological processes. Finally, a generic process of distribution of matter and energy ensures homogeneity within the ecosystem. The formalism of signed digraphs (Levins, 1974) is employed in Figure 2B, emphasizing classical feedbacks as positive (the arrow) or negative (the solid dot) between compartments.

The minimum ecosystem defined as such, requires four variables: R, which represents the state of the nutrient pool, P, the state of the primary production population, M, the state of the pool of detrital organic matter, and D, the state of the decomposer population, and assumes that the units are all the same. The model is formulated by a system of four ordinary differential equations as:

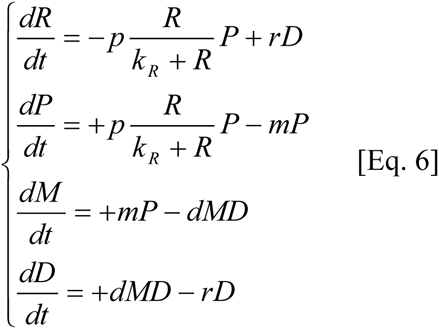

where *p* is a production rate (time^-1^), *r*, a remineralization rate (time^-1^), *m*, a primary producers mortality rate (time^-1^), and *d*, a decomposition rate (unit of state^-1^.time^-1^). The constant, k_R_ (units of R) is a half-saturation constant of the Holling type II function (Holling, 1959) that regulates intake of nutrients by primary producers. The ecosystem is conservative in terms of matter; the sum or derivatives are equal to zero, hence R+P+M+D = I_0_.

We then fix a set of initial conditions 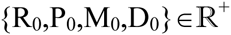 which are the supposedly known conditions at time t_0_. Equilibriums were calculated when time derivatives are all equal to zero [Eq. 7], and their stability properties are determined by studying the sign of the derivative around the calculated solutions:

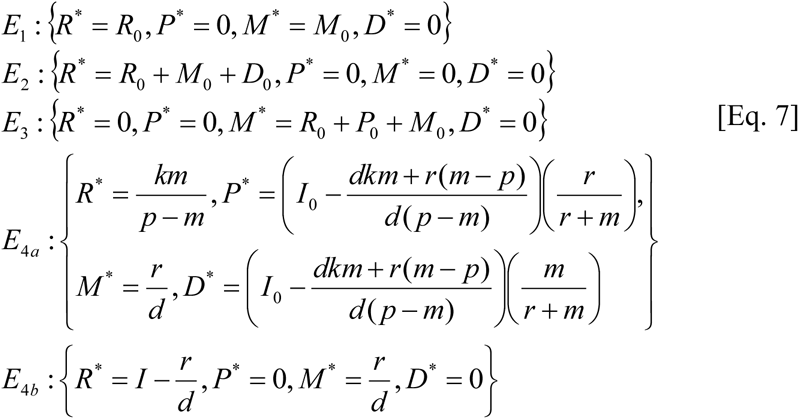

where R_0_ > 0, P_0_ > 0, M_0_ ≥ 0 and D_0_ >0, and *a fortiori* I_0_ = R_0_ + P_0_ + M_0_ + D_0_ > 0. All five equilibriums listed above are stable and coexisting with the unstable trivial equilibrium {R^*^=0, P^*^=0, M^*^=0, D^*^=0}. E_4a_ is reached if p > m and E_4b_ is reached otherwise (assuming that the decomposers are acting fast with respect to the dynamics of the entire system). E_1_, E_2_ and E_3_ equilibriums do not respect our definition of an ecosystem:

- E_1_ is the case of no living organisms at the beginning (spontaneous generation is not allowed), and
- E_2_ and E_3_ are equilibriums with the initial absence of the primary producer or decomposer populations respectively, leading to the extinction of the other population (hence the condition of the co-existence of P and D is not fulfilled).

### Calculating changes in receptors and modelling the influence of drivers of change

In the model presented above, many receptor variables X can be identified. They can be the state variables (mainly representing the living populations, *i.e.* P or D) or the processes (like the ecosystem functions: primary production, decomposition and nutrient recycling). For all these variables, we calculated an impact as δ = Δ/X^*^, the relative variation from the baseline X^*^, consecutive to a virtual project implementation. *D* is the difference between two equilibrium values X^*^ to X^**^, after a change in states (such as nutrient or detrital organic matter inputs) or parameters (mostly decreases in primary production rate, increases in primary producers’ mortality rate, decrease in decomposition and recycling rates) consecutive to project implementation.

For the Environmental Impact Assessment, it is only required to know the amplitude of the changes consecutive to modifications of states or parameters to predict an impact on receptors. However, since we wish to include socio-economic aspects, we linked in a second step the change in ecosystem state and function to the possible influence of stakeholders on the project development (or the project ‘Life Cycle’). The project development is controlled by groups of stakeholders, and the related "activity" depends on many factors that do not depend directly on ecosystem feedbacks (Binder *et al.*, 2013).

Treating a ‘socio-economic-ecological system’ using systemic principles generates outcomes with little interest due to possible socio-economic feedbacks that are not connected as reactions to a physical system (*i.e.* "A" has an action on "B", and in return, "B" modifies "A", as in Figure 2B). We thus revise the notion of feedbacks by "A" has an action on "B" until "A" <u>*realizes that*</u> the action on "B" can be unfavourable to its own development. This formulation partly overlaps with the notion of “vulnerability” presented in Toro *et al.*, 2012 and “risk” (Gray and Wiedemann, 1999). The socio-economic system is introduced as a driver of change for the minimum ecosystem, instead of as a state variable like in other SES frameworks (Binder *et al.*, 2013).

Consequences for the impacts on receptors are described in terms of the relative "activity" A (A *Î*[0 1]) of the project, related to the change in states or parameters by minimal linear functions (*i.e.* if *x* represents any potential change rates - in parameters or states - the effective change rates, *y*, are expressed by *y* = A*x*). The project activity is calculated as the complement of the relative socio-economic cost, C, of project development, expressed as:

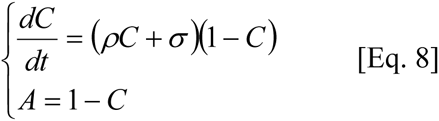

where σ is a relative social awareness rate (increase, in time^-1^, of the number of stakeholders aware of the negative consequences of the project within the total number of stakeholders), and *r* is the reactivity rate (the standardized speed, in time^-1^, at which the socio-economic cost corresponding to mitigation or remediation measures increases).

All simulations and related calculations were performed using open source software (Scilab Enterprises, 2012).

## RESULTS

### Examples of the impact predictions estimated by the model

Three different scenarios were set-up for specific receptors (Table 1). Examining the steady-states of the system and their stability stresses the position of the set of parameters θ = {p, m, d, r, k_R_}and their relative importance in the definition of the system equilibrium. For building scenarios, it is assumed that the parameters’ orders of magnitude are:

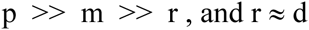

**Table 1.**
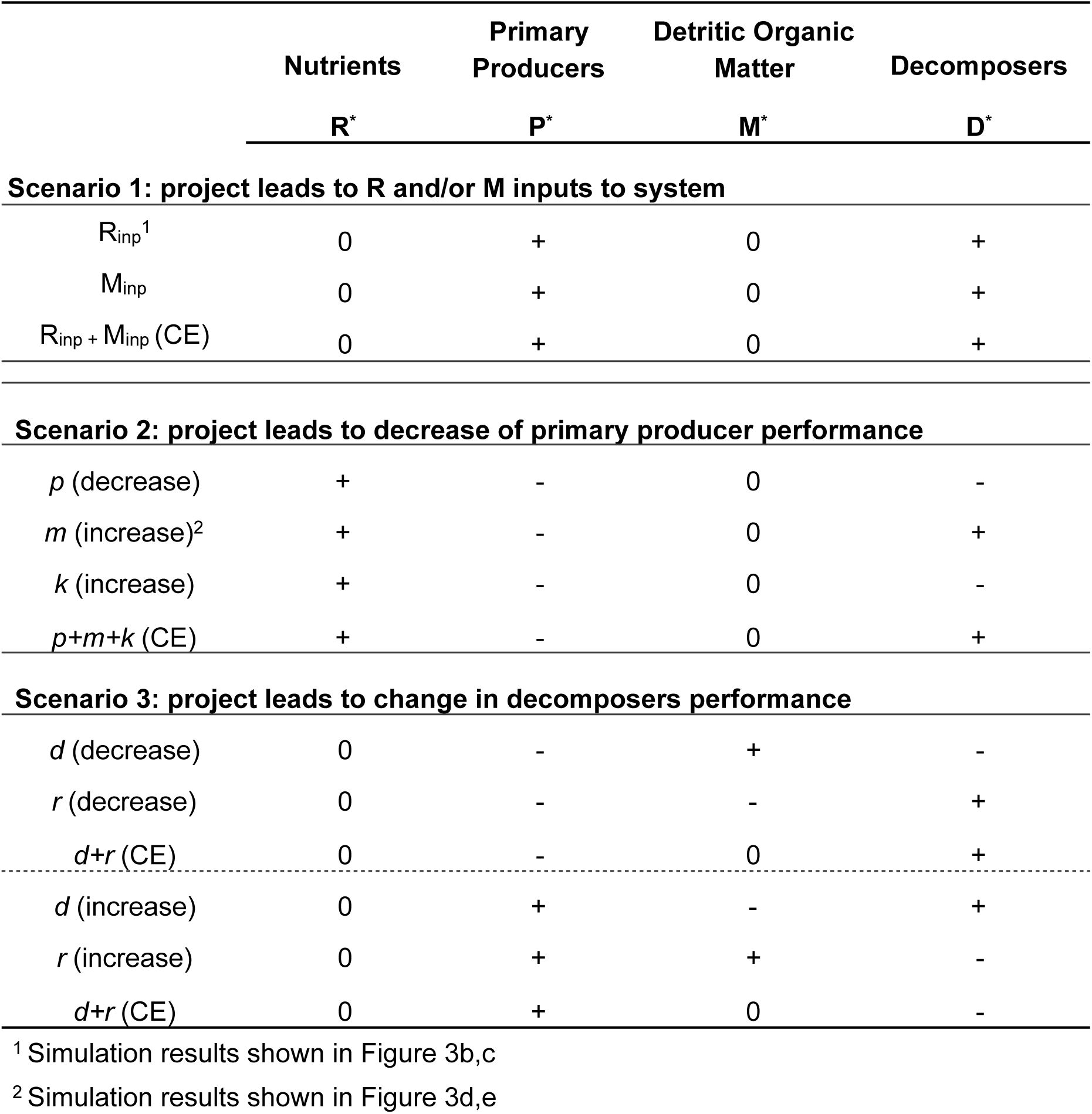
Summary of model outcomes for three scenarios. Relative changes in impact are calculated in terms of mass or energy content and compared for the scenarios described in the results.

|  | Nutrients | Primary Producers | Detritic Organic Matter | Decomposers |
| --- | --- | --- | --- | --- |
|  | R* | P* | M* | D* |
| **Scenario 1: project leads to R and/or M inputs to system** |  |  |  |  |
| $R_{inp}^1$ | 0 | + | 0 | + |
| $M_{inp}$ | 0 | + | 0 | + |
| $R_{inp} + M_{inp}$ (CE) | 0 | + | 0 | + |
| **Scenario 2: project leads to decrease of primary producer performance** |  |  |  |  |
| $p$ (decrease) | + | - | 0 | - |
| $m$ (increase)<sup>2</sup> | + | - | 0 | + |
| $k$ (increase) | + | - | 0 | - |
| $p+m+k$ (CE) | + | - | 0 | + |
| **Scenario 3: project leads to change in decomposers performance** |  |  |  |  |
| $d$ (decrease) | 0 | - | + | - |
| $r$ (decrease) | 0 | - | - | + |
| $d+r$ (CE) | 0 | - | 0 | + |
| $d$ (increase) | 0 | + | - | + |
| $r$ (increase) | 0 | + | + | - |
| $d+r$ (CE) | 0 | + | 0 | - |

<sup>1</sup> Simulation results shown in Figure 3b,c<sup>2</sup> Simulation results shown in Figure 3d,e

Nonetheless, d is controlled by the quantity of substrate available. k is considered as small and the primary producers being assumed to have a good affinity for the available nutrients. When changes of parameters were simulated (as in Scenarios 2 and 3) they were varied in the same proportions. Inputs were simulated separately and then cumulated (CE), and their impacts on the 4 state variables at equilibrium (R^*^, P^*^, M^*^ and D^*^) were examined.

The first scenario simulated direct inputs of nutrients and detrital organic matter. Results show that in all cases, R^*^ and M^*^ did not vary (despite their initial increase). On the contrary, the variables representing living compartments, P^*^ and D^*^, increased. Results also show that the relative variation to the baseline, *d*, is identical for P^*^ and D^*^ (both positive deviations, Table 1). Concerning processes at equilibrium, the primary production and the primary producer mortality both increased, as well as the processes of decomposition and recycling, since none of these parameters were affected by the project implementation.

The second scenario simulates an impact which consists of the decrease in primary producer performance. This could be due to the physiological capacities of the organisms being affected by the project or because the environmental conditions limit their expression (*e.g.* a strong increase in water column turbidity). In this situation, the parameters affected are k and m (which increased), and p (which decreased). It should be recalled that p was kept greater than m (p - m > 0), as per our parameter hierarchy. A decrease of p and an increase of k (global decrease of primary productivity) always has a negative effect on P^*^ (hence on primary production), a positive effect on R^*^, and a negative effect on D^*^. In both cases, the relative variations to the baseline, *d*, are identical for P^*^ and D^*^. An increase of m has a similar effect on P^*^ and R^*^, but has a negative effect on D^*^. The cumulative effect (p + m + k) is almost equal in magnitude to the effect of a decrease in m, which is much higher (by several orders of magnitude) than the effects of p and k. Effects of p and k are quite negligible, each having a typical order of magnitude of the parameters in θ.

The third and final scenario simulated a change in the decomposer activity. This could be triggered by a change in taxonomic composition, and also by the action of chemical substances released during the project. Decreases and increases in d and r were simulated, first separately and then together. Changes in d and r have no effect on R^*^. A decrease of d as a negative effect on P^*^ (hence decreasing primary production) and D^*^, and logically, an increase of d has a positive effect on P^*^ (thus the increasing primary production) as well as D^*^. In both cases, the relative variations to the baseline, δ, are identical for P^*^ and D^*^. Effects of a decrease or an increase in r on P^*^ and D^*^ are opposed. P^*^ increases and D^*^ decreases when r increases, and P^*^ decreases and D^*^ increases when r decreases. Cumulative effects reinforce slightly the effect of a change in r which is largely predominant in the dynamics of P and D. The changes of d and r affect the primary production *via* a change in the availability of R. When the recycling is enhanced (mainly by the increase of r but also by an increase of d), R production increases but an excess of R is used to increase the state of the primary producer P. It is because the production rate p is high compared to r, that R* is not affected by changes in r or d. Changes in r and d have opposite effects on M^*^. A decrease (respectively, increase) of d has a positive (respectively, negative) effect on M^*^, and a single decrease (respectively, increase) of r has a negative (respectively, positive) effect on M^*^. When changes are cumulated (in equal proportions), the effect of changes in r and d on M^*^ is null, showing that they have the same amplitude on M^*^.

### Behaviour of system when drivers of change were included

In the impact assessment *per se*, the effects of changes in ecosystems components (states and functions) were considered as a deviation of stable equilibrium values regardless of the time scales of the transitory phase. The consequences of introducing socio-economic drivers were considered by numerical simulations. To take into account the potential influence of socio-economic drivers, simulations were performed introducing explicitly a changing rate that depends on the relative project activity within Equation 6, affecting either states or parameters. Figure 3 shows results of simulations for just two different examples of impact taken from Table 1. The first scenario illustrated (Figure 3b, c) is for a project development that induces a change in state (a nutrient input triggering an initial increase of R, scenario 1), and the second illustration (Figure 3d, e) suggests what can occur when a project induces a change in parameters (in this case an increase in the mortality rate of primary producers and hence a decrease of their survival, scenario 2). The reactivity rate *r* was set to 0.02 (time^-1^) and the awareness rate *s* was set to 10^-4^ (time^-1^). For both scenarios, the project activity starts at t = 200 (time), the dynamics being considered at steady state before. Figure 3a shows the activity of the project reaches instantaneously 1 at ‘time’ 200 when the project is implemented and then decreases smoothly as global awareness of negative impacts among stakeholders’ increases [Eq. 8]. The project activity thus decreases to 0 by ‘time’ 800. This is a consequence of the relative socio-economic cost of the project reaching 1, which in our model, defines the limit of the exploitability of the project (*i.e.* when all possible time and resources are being invested in side issues).

**Figure 3.**
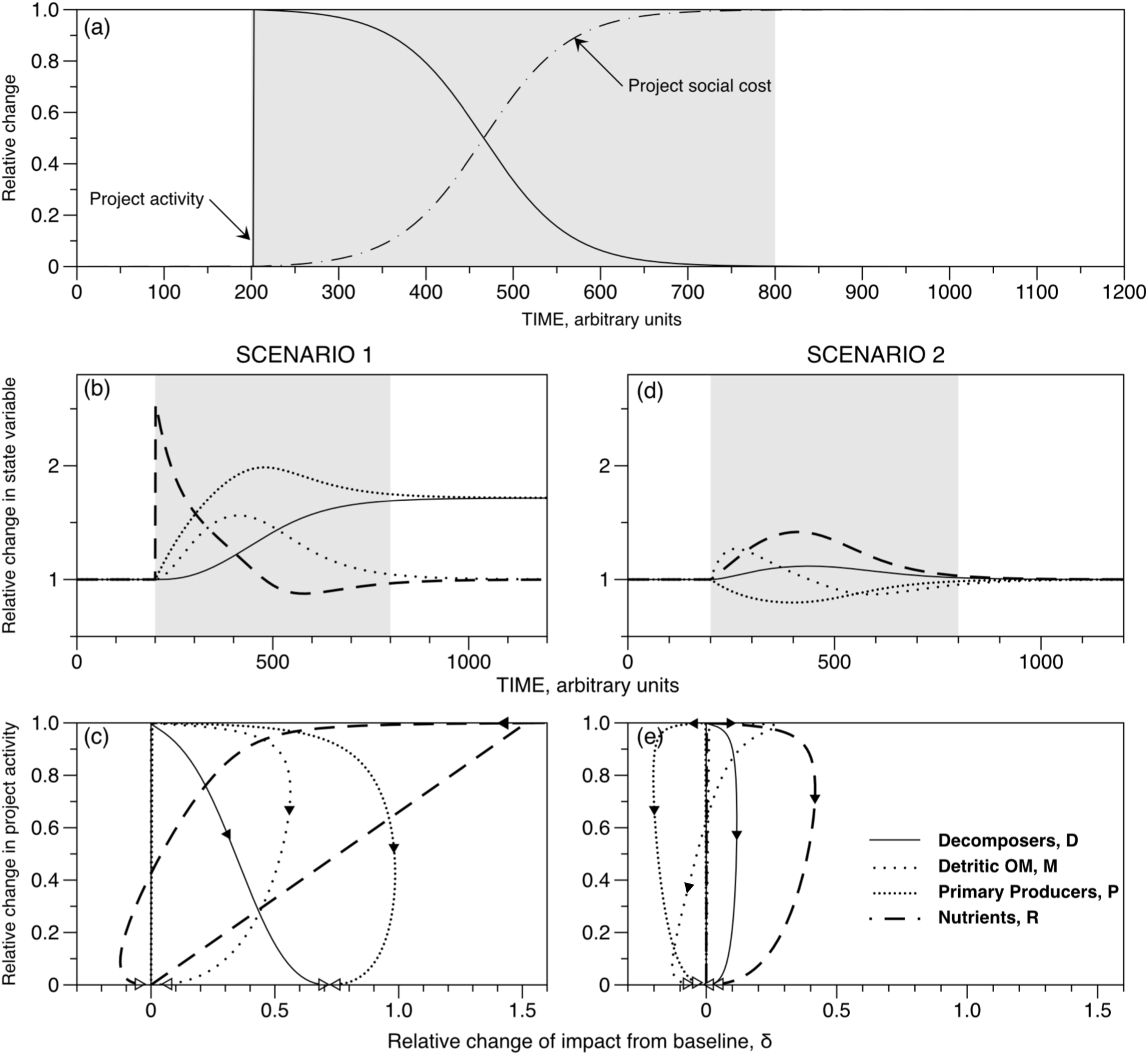
Impact as influenced by stakeholder awareness and project cost-effectiveness. (a) Inverse relationship between the Project Activity and Project social cost (awareness of a negative impact among stakeholders) for the simulated scenarios. The grey shaded area is the project activity duration (between time step 200 and 800 here). Behaviour of the four state variables (b, d) and the relative changes in impact (c, e) during scenario 1 and 2, respectively. These scenarios are also listed in Table 1. Filled triangles indicate in which direction the relative impacts are changing for each of the four compartments as the state variables evolve (b, d), and the unfilled triangles are placed at or near the end of the curves. All curves start at “0” in these simulations.

In the first scenario, when R increased sharply, both P and D increased as well, but more slowly (Figure 3b). When the project activity stopped (outside the grey area, after ‘time’ 800), all states have reached an equilibrium, which is, for M, the equilibrium prior to the implementation of the project, but for P and D, a different higher equilibrium. In that sense, the outcome is similar to the outcome of the previous scenario 1. Figure 3c shows that the δ for P and D varies differently showing the modulation by the project activity tends to alter the final amplitude of the impacts on each of the receptors.

In the second scenario, the configurations for the relative socio-economic cost and activity of the project are identical, but the outcomes were very different from those in scenario 2. In this case, when project activity stopped, causes for changes in the mortality rates disappeared and equilibrium states came back to the values prior to the project implementation (Figures 3 d, e). Therefore, around ‘time’ 400, the impact of the project on all receptors reaches a maximum, but all impacts relative to the baseline, δ, decreased and returned to zero afterward (Figure 3e).

## DISCUSSION

The practice of EIA arose from a societal imperative to have documented expertise about potential impacts on the environment from development projects. This was the result of a legal framework created to defend environmental quality of communities and regions in the US (Cashmore, 2004; Morgan, 2012), and coinciding with a rise in visibility of ecological sciences (Supplementary Information, Figure A). Subsequently, similar requirements for environmental impact assessment were adopted by a majority of countries (Morgan, 2012). This has engendered repeated calls to develop a theory of impact assessment (Lawrence, 1997) as the practice dispersed. The need for an EIA process created a profession with a vital role in the safeguard of environmental quality, but that relies heavily on disputable methods and has an uneven record (*e.g.* Wood, 2008; Wärnbäck and Hilding-Rydevik, 2009; Barker and Jones, 2013). Public pressure from stakeholders may provide some measure of accountability, however, *post hoc* analyses are rare (Lawrence, 1997) and systems can differ significantly between countries (Lyhne *et al.*, 2015). Critical review may only happen in the aftermath of a dramatic accident, such as the Macondo well blow-out in 2010 (US Chemical Safety and Hazard Investigation Board, 2016) or after management failures (Rotherham *et al.*, 2011).

### The value of quantification

Our study reflects on the two main scientific components of EIAs: expertise and prediction. The first is the role of the expertise. We have stressed the needs for the experts to identify receptors and to provide proper estimates of baselines. The second one is the ability of ecological theory to prediction ecosystem dynamics. We have emphasized the critical importance of the formulation of the ecosystem model to calculate correctly baselines and predict impacts. The intention of Leopold *et al.* (1971) was however far from this approach. Their approach consisted in providing a sort of template for EIA and EIS documents and to ensure a common logic for how the “magnitude and importance” of the impacts identified would be presented to federal evaluators. They did not provide any details about how exactly impacts would be assessed beyond a comparison between conditions before and after the project. We therefore replaced this generic matrix approach by a quantification of system dynamics, which allows scenarios to be designed and tested.

### Receptor selection

Scenarios are selection of the possible combinations that could be examined, and which are usually specific to the type of project that would be implemented. The ecosystem model is then used as a tool to helps experts identifying specific receptors. Receptors can only be identified if their *δ* is different from zero (either strictly positive or strictly negative). It can be identified easily in Table 1, but this is not the only condition: to be a receptor the *δ* must indeed be greater (in absolute value) than the *δ*\* corresponding to the limit of detection of the impact [Eq 5]; this is a statistical concept required to estimate the dispersion of the values of the receptor variables around their average. These two conditions then define what receptors are. Receptors are indeed subject to change and must be sensitive enough to be detectable with the statistical tests applied. Hence, an EIA, in contrast with a risk assessment, implies automatically a change in the receptors and aims to quantify them with a defined level of certainty and accuracy. A consequence of this is that if two receptor variables were identified as having the same dispersion, the impact will be better assessed if the averages have higher values. For example, in a marine system, the biomass of decomposers D, can be much greater than the biomass of the primary producers, P (Simon *et al.*, 1992), which means that it could be better to assess impact on D, than on P. This can be completely different for terrestrial ecosystems (Cebrian and Lartigue, 2004).

### Baselines and reference conditions

In our model, the description of changes is based on the calculation of equilibrium (the baseline) and their stability, and then follows the displacement of the equilibrium values under changes in state variables, forcing variables, or parameters (Figure 3b-e). This description is a basis for clarifying our understanding of the problem. A dynamic model constrains our investigation to plausible causal relationships between the variables (receptors) and permits us to explore their contribution to the entire system. The dynamic behaviour provides a point of reference for comparisons between scenarios (as shown in Table 1 and Figure 3), or as they correspond to a specific project development. Formulating a minimum ecosystem as an example, has shown that complex behaviours can emerge with only four state variables. These results illustrate for the first time the dynamics of impact responses by receptors, revealing how complicated the evaluation of recommendations to mitigate impact may be. Furthermore, this underscores the importance of monitoring to ensure accountability over the project life cycle, including cumulative effects.

### Minimum ecosystems and complexity

Models are simulation tools which aid exploration of possible outcomes and the evaluation of the simulated baseline, as well as the relevance to simulated scenarios (Tett *et al.*, 2011). Our minimum ecosystem model is essentially a representation of a perfect and autonomous bioreactor, which does not exist, nor can one be created as presented. Nutrients and detrital organic matter are 100% recyclable by one functional group of decomposers. Populations are stable indefinitely if conditions on the parameters (essentially p > m) are respected. These conditions are not realistic, but serve the development and presentation of our approach. The proposed procedures can be applied to more complex systems, encompassing large quantity of variables (or compartments) as well as non-linear processes and hybrid dynamics, like what would be expected in more realistic representations of ecosystems.

However, the condition that a certain form of stability can exist in the system must be respected. It should be noted that the question of stability in ecology is part of an on-going scientific discussion recently summarized by Jacquet *et al.* (2016). This is critical to environmental impact theory because it is the presumption of stability which ensures the baseline is maintained (does not drift) during the project life cycle (Thorne and Thomas, 2008; Pearson *et al.*, 2012). In other words, an EIA is supposed to certify that what is measured as change only corresponds to an impact from the project, not external variations. Hence, monitoring takes on a new importance. For example, monitoring a non-impacted site as a reference to detect possible ecosystem drift, may be one way to assure that this condition of baseline stability is valid. This solution is conditioned itself by the necessity to have a reference site which can be characterized by exactly the same ecosystem.

The second basic assumption of our minimum ecosystem implies that the distribution of elements is homogeneous inside the project area. This is not always (and even rarely) the case and in aquatic systems, hydrodynamics leads to partial mixing that cannot be assimilated to complete homogeneity. Therefore, accounting for the spatial distribution structure of the elements would require the model structure be modified. For example, we can use partial differential equations or any other formulation that can treat spatial covariance. When spatial covariance is proven to exist for relevant receptors, the corresponding statistics for the test of impact must account for the spatial covariance using geostatistical methods (*e.g.* Agbayani *et al.*, 2015; Wanderer and Herle, 2015).

### Socio-ecological systems

The idea that all components (*i.e.* Environmental, Social, Health … impacts) can be inserted into a single system framework remains quite challenging. While a considerable number of propositions for conceptual frameworks and planning charts exist (Haberl *et al.*, 2009; Binder *et al.*, 2013; Bowd *et al.*, 2015; Ford *et al.*, 2015) offering some insights into the complex social interactions and policy constraints involved, there is little in the way of theoretical development for impact theory. We only studied here the project activity controlled by its socio-economic cost (side costs being related to remediation and mitigation measures) as a driver of ecosystem changes. We have not, for example, considered that changes in some receptors can trigger an increase in cost and a decrease in activity. In other words, we have not considered feedbacks between the receptors and cost, because it did not appear clearly how awareness of stakeholders and reactivity of managers could be directly linked to changes in receptors (Binder *et al.*, 2013; Bowd *et al.*, 2015) for which “acceptable” remediation or mitigation measures should have already been considered during the process (Figure 2B; Drayson and Thompson, 2013). Indeed, stakeholders’ awareness depends on many factors, like information or education (Zobrist *et al.*, 2009), and reactivity of managers can be constraints by many other economic and political factors (Ford *et al.*, 2015). However, the minimal model that we proposed for expressing the dynamics of the drivers of change [Equation 6] can (and should) become more rich to take into account more complete descriptions of the mechanisms that modulate awareness, activity and reactivity rates within sociological networks. We suggest that our approach could be particularly useful in the scoping step as a means to explore possible scenarios outcomes.

## CONCLUSIONS

This study has linked statistical tests and mathematical modelling to assess an impact and consider some of the socio-economic drivers that mitigate it. This constitutes a first step toward an ecosystem-based approach for EIA, which needs to be proven and improved. If technically, there are possibilities for EIA to rest on objective quantitative approaches, these can only be valid if the predictive capacity of the model is assured. This was, and still is, a major limitation. Furthermore, all forms of environmental impact assessment are complicated by the absence of fundamental laws in ecology (Lange, 2002) which has limited the understanding of complex objects in ecosystems. Most of the time, ecosystem models simulate dynamics with properties that are not found in realistic systems (May, 1977). We believe that to progress toward quantitative EIA it is necessary to build much closer, interdisciplinary collaborations between applied and fundamental research on ecosystems, to overcome the historical divergences. This exchange could be encouraged through concrete measures such as including funding for fundamental development within EIA as well as requiring that data collected for IA be made available in open source repositories, accessible for fundamental research.

## SUPPLMENTARY MATERIAL

This material is not included in this version.

## ACKNOWLEDGEMENTS

This work was presented in part at the ICES meeting “MSEAS2016” in Brest, France and is part of the dissertation of J Coston-Guarini, inspired by the original insights of Jody Edmunds.

